# The rice OsERF101 transcription factor regulates the NLR Xa1-mediated perception of TAL effectors and Xa1-mediated immunity

**DOI:** 10.1101/2021.11.12.468346

**Authors:** Ayaka Yoshihisa, Satomi Yoshimura, Motoki Shimizu, Sayaka Sato, Akira Mine, Koji Yamaguchi, Tsutomu Kawasaki

## Abstract

- Plant nucleotide-binding leucine-rich repeat receptors (NLRs) initiate immune responses and the hypersensitive response by recognizing pathogen effectors. *Xa1* encodes an NLR with an N-terminal BED domain, and recognizes transcription activator-like (TAL) effectors of *Xanthomonas oryzae* pv. *oryzae* (*Xoo*). The molecular mechanisms controlling the recognition of TAL effectors by Xa1 and the subsequent induction of immunity remain poorly understood.
- Xa1 interacts in the nucleus with two TAL effectors via the BED domain. We identified the AP2/ERF-type transcription factor OsERF101/OsRAP2.6 as an interactor with Xa1, and found that it also interacts with the TAL effectors. Overexpression of *OsERF101* exhibited an enhanced resistance to an incompatible *Xoo* strain only in the presence of Xa1, indicating that OsERF101 functions as a positive regulator of Xa1-mediated immunity. Unexpectedly, *oserf101* mutants also showed enhanced Xa1-dependent resistance, but in a different manner from the overexpressing plants. This result revealed an additional Xa1-mediated immune pathway that is negatively regulated by OsERF101. Furthermore, OsERF101 directly interacted with the TAL effectors.
- Our results show that OsERF101 regulates the recognition of TAL effectors and the Xa1-mediated activation of the immune response. These data provide new insights into the molecular mechanism of NLR-mediated immunity in plants.

## Introduction

Plants have developed two tiers of immune systems to defend against pathogen infection. The first tier is initiated through recognition of microbe-associated molecular patterns by plasma membrane-localized pattern recognition receptors. This is referred to as pattern triggered immunity (Dangl *et al.*, 2013). To inhibit pattern triggered immunity or to improve the nutrient conditions suitable for pathogen proliferation, pathogens deliver a variety of effectors into plant cells (Dou & Zhou, 2012). For the second tier of immune responses, plants have evolved a family of <u>n</u>ucleotide-binding (NB) leucine-rich repeat (LRR) receptors (NLRs) that directly or indirectly recognize pathogen effectors (Jones *et al.*, 2016). This is referred to as effector triggered immunity. Effector triggered immunity often involves a hypersensitive cell death response (HR). Several NLRs indirectly recognize pathogen effectors by interacting with host factors. These host factors are defined as “sensing decoys” that mimic the targets of pathogen effectors and act as co-receptors with the NLRs (Paulus & van der Hoorn, 2018).

*Xanthomonas oryzae* pv. *oryzae* (*Xoo*) causes rice bacterial blight disease, one of the most important rice diseases in the world. *Xoo* has developed transcription-activator like (TAL) effectors to facilitate bacterial growth. TAL effectors contain a central region of polymorphic repeats (central repeat region (CRR)) with each repeat consisting of 34 amino residues, several nuclear localization signals (NLSs), and an activation domain at the C terminus. Each of the repeats in the CRR specifies a nucleotide for binding (Boch *et al.*, 2009; Moscou & Bogdanove, 2009). *Xoo* secrets TAL effectors into rice cells through a type III secretion system, and then the TAL effectors localize to the host nuclei in order to regulate expression of certain host genes. One of the *Xoo* TAL effectors, AvrXa7, accelerates expression of the *SWEET14* gene, which encodes a plasma membrane-localized sugar transporter (Antony *et al.*, 2010). The enhanced accumulation of SWEET14 protein increases the efflux of sugars from the cytoplasm to the apoplast, and this provides additional nutrients for the pathogen (Naseem *et al.*, 2017). This process is very important for bacterial virulence, because defects in the expression of *SWEET14* greatly reduce bacterial growth (Oliva *et al.*, 2019).

The bacterial blight disease resistance gene *Xa1* was identified originally in the rice cultivar Kogyoku (Yoshimura *et al.*, 1998). It encodes an NLR protein with an N-terminal BED-type zinc finger domain. Xa1 recognizes TAL effectors and induces immune responses including the HR (Ji *et al.*, 2016), but the mechanisms for these processes have not been elucidated. Recently, alleles of *Xa1* called *Xa2*, *Xa14*, *Xa45*, and *Xo1* have been isolated (Ji *et al.*, 2020; Read *et al.*, 2020; Zhang *et al.*, 2020). Their predicted protein structures indicate that the BED and NB regions are highly conserved, but their C-terminal LRR regions are distinguished by the number of repeats. This suggests that the LRR regions may determine the interactions with TAL effectors (Read *et al.*, 2020).

The immune responses induced when Xa1 and its allelic NLR proteins recognize TAL effectors can be suppressed by interfering TAL (iTAL) effectors, also referred to as truncated TAL (truncTAL) effectors (Ji *et al.*, 2016; Read *et al.*, 2016; Ji *et al.*, 2020). iTAL effectors lack the activation domain but retain the NLSs. iTAL effectors can be classified into two types (A and B) based on their structures (Ji *et al.*, 2016; Read *et al.*, 2016). However, the molecular mechanisms by which iTAL effectors suppress Xa1-mediated immunity remain to be revealed.

Rice contains 139 AP2/ERF-type transcription factors (Nakano *et al.*, 2006). The ERF transcription factors are involved in a variety of cellular processes including development and responses to biotic and abiotic factors. The OsERF101/OsRAP2.6 transcription factor has been reported to participate in resistance to rice blast (Wamaitha *et al.*, 2012), leaf senescence (Lim *et al.*, 2020), and drought stress (Jin *et al.*, 2018).

In this study, we found that the BED domain of Xa1 forms a complex with two TAL effectors, AvrXa7 and Xoo1132. To understand the molecular mechanism of Xa1-mediated immunity, we screened for proteins that interact with Xa1, and identified OsERF101. We investigated the interactions between OsERF101 and Xa1 in plants. We also analyzed Xa1-dependent resistance and TAL effector-induced gene expression using plants that overexpressed *OsERF101* and plants carrying knockout mutations of *OsERF101*. Furthermore, we found that OsERF101 directly interacts with the TAL effectors. The results of these experiments suggest that OsERF101 is likely involved in both effector recognition and immune activation mediated by Xa1.

## Materials and Methods

### Plant materials

Rice (*Oryza sativa*) *Japonicum* cultivars Kogyoku and Nipponbare were used as the wild-type plants. The *xa1* and *oserf101* mutants were generated using the CRISPR/Cas9 system as described below. The *OsERF101*-OX plants were generated as described below.

### Plant Transformation

To construct the plasmids for the CRISPR/Cas9 system, we used the guide RNA cloning vector pU6gRNA and the all-in-one Cas9/gRNA vector pZDgRNA_Cas9ver.2_HPT, which were kindly provided by Dr. Masaki Endo (Mikami *et al.*, 2015). The 20 bp sequences from 108 to +127 of *Xa1* (5’-GCAACTGGTCTGCAAAGATC-3’) and from 206 to +225 of *OsERF101* (5’-GTGTTCGACAGCGGCCATGG-3’) were selected as the target sites of Cas9 by using the CRISPR-P website (http://cbi.hzau.edu.cn/cgi-bin/CRISPR). These DNA fragments were cloned into pU6gRNA, and then subcloned into pZDgRNA_Cas9ver.2_HPT (Mikami *et al.*, 2015). Calli generated from rice embryos were transformed using *Agrobacterium tumefaciens* EHA101 carrying each construct, as described previously (Hiei *et al.*, 1994). To confirm the mutations, the genomic regions containing the Cas9 target sites were amplified by PCR and sequenced as previously described (Mikami *et al.*, 2015). For overexpression, the entire coding region of *OsERF101* was amplified with gene specific primers (Table S1) using cDNAs prepared from Kogyoku leaves as the templates, and the PCR product was cloned into the binary vector pGWB2 (Nakagawa *et al.*, 2007). In the resulting construct, the *OsERF101* gene was driven by the CaMV 35S promoter. Calli generated from rice embryos were transformed with the construct as described previously (Hiei *et al.*, 1994).

### Transient assays using rice protoplasts

Protoplasts were isolated from cultured rice cells by digestion of the cell walls with Cellulase RS (Yakult) and Macerozyme R-10 (Yakult) as described previously (Yamaguchi *et al.*, 2013). Aliquots (100 μl) of protoplasts (2.5 × 10^6^ cells / ml) were transformed with plasmid DNA using the polyethylene glycol (PEG) method (Chen *et al.*, 2010). For subcellular localization analysis and BiFC analysis, transfected protoplasts were observed using a fluorescence microscope, the Axio Imager M2 (Carl Zeiss) with the ApoTome2 system (Carl Zeiss). The BiFC analyses were carries out as reported previously (Yamaguchi *et al.*, 2013).

### Split NanoLuc Luciferase complementation assay

DNA fragments of *Xa1*, *BED*, *NB*, *LRR*, *AvrXa7*, and *Xoo1132* were transferred into *p35S-LgBiT-T7-GW* or *p35S-SmBiT-T7-GW* using the Gateway system with LR clonase reactions (Taoka *et al.*, 2021). The Firefly *Luciferase* gene controlled by the CaMV 35S promoter was used as an internal control. The constructs were transfected into rice protoplasts as described above. After 18 h incubation at 30°C, the activities of the Firefly and NanoLuc luciferases were measured in a TriStar2 LB942 luminometer (Berthold) using the ONE-Glo Luciferase Assay System (Promega) and the Nano-Glo Live Cell Assay System (Promega).

### Yeast two-hybrid assays

The yeast two-hybrid screening and interaction assays were based on the requirement for histidine for yeast growth, as described previously (Ishikawa *et al.*, 2014).

### Protein extraction and immunoblotting

Total proteins were extracted from rice protoplasts in a buffer including 50 mM Tris-HCl pH 7.5, 1 mM EDTA, and a protease inhibitor cocktail (Roche), and analyzed by immunoblotting with α-HA, α-Lg, and α-T7.

### Pathology Assays

Fully expanded rice leaves were inoculated with *Xoo* T7174 or *Xoo* T7133 by infiltration of the bacterial suspension (OD_600_ = 0.25) with a needleless syringe. The bacterial populations in the leaves after infiltration were analyzed by quantitative realtime PCR. The DNA levels of the *Xoo XopA* gene relative to those of the rice *ubiquitin* gene were measured using genomic DNAs purified from the infected leaves.

### RNA isolation and quantitative real time PCR

Total RNA was isolated from rice leaves using TRIzol reagent (Invitrogen) and then treated with RNase-free DNase I (Roche). First-strand cDNA was synthesized from 1 μg total RNA using an oligo-dT primer and ReverTra Ace reverse transcriptase (Toyobo). Expression levels were measured by quantitative real time PCR using the SYBR Green master mix (Applied Biosystems) in a Step-One Plus Real-Time PCR system (Applied Biosystems). The expression levels were normalized against a *ubiquitin* reference gene. Three biological replicates were used for each experiment, and two quantitative replicates were performed for each biological replicate.

### RNA-seq

Total RNA was used to make sequencing libraries using a NEBNext Ultra RNA Library Prep Kit for Illumina (NEB, USA) according to the manufacturer’s instructions. Subsequently, the libraries were sequenced using an Illumina Hiseq4000 platform to obtain 150 bp paired-end reads. FaQCs (Lo & Chain, 2014) software was used to filter high quality reads from the generated sequence reads. The filtered reads were aligned with the rice reference sequence for Nipponbare (http://rice.plantbiology.msu.edu/pub/data/Eukaryotic_Projects/o_sativa/annotation_dbs/pseudomolecules/version_7.0/all.dir/all.con) using Hisat2 (Kim *et al.*, 2015) software, and counted for each gene using FeatureCounts (Liao *et al.*, 2014) software. DEGs were detected using a TCC package (Sun *et al.*, 2013).

## Results

### Xa1 interacts with TAL effectors in the rice nucleus

In order to analyze the interactions between Xa1 and TAL effectors, we first used Xa1 protein fusions with green fluorescence protein (GFP) to determine their subcellular localization in transiently expressing rice protoplasts. GFP was fused to the N terminus of full length Xa1^1-1,802^ and four different Xa1 regions, Xa1^1-325^, Xa1^1-1,012^, Xa1^312-1,012^, and Xa1^1,008-1,802^. These four regions contained the BED, BED-NB, NB, and LRR domains, respectively (Fig. 1a). GFP-Xa1^1-325^ and GFP-Xa1^1-1,012^ were localized mainly in the nucleus (Fig. 1b,c). GFP-Xa1^1-1,802^ was detected in both the nucleus and the cytoplasm, while GFP-Xa1^312-1,012^ and GFP-Xa1^1,008-1,802^ were localized only in cytoplasm. Thus, it is likely that BED possesses a nuclear-localization activity, which is consistent with a recent report (Zhang *et al.*, 2020).

**Fig. 1.**
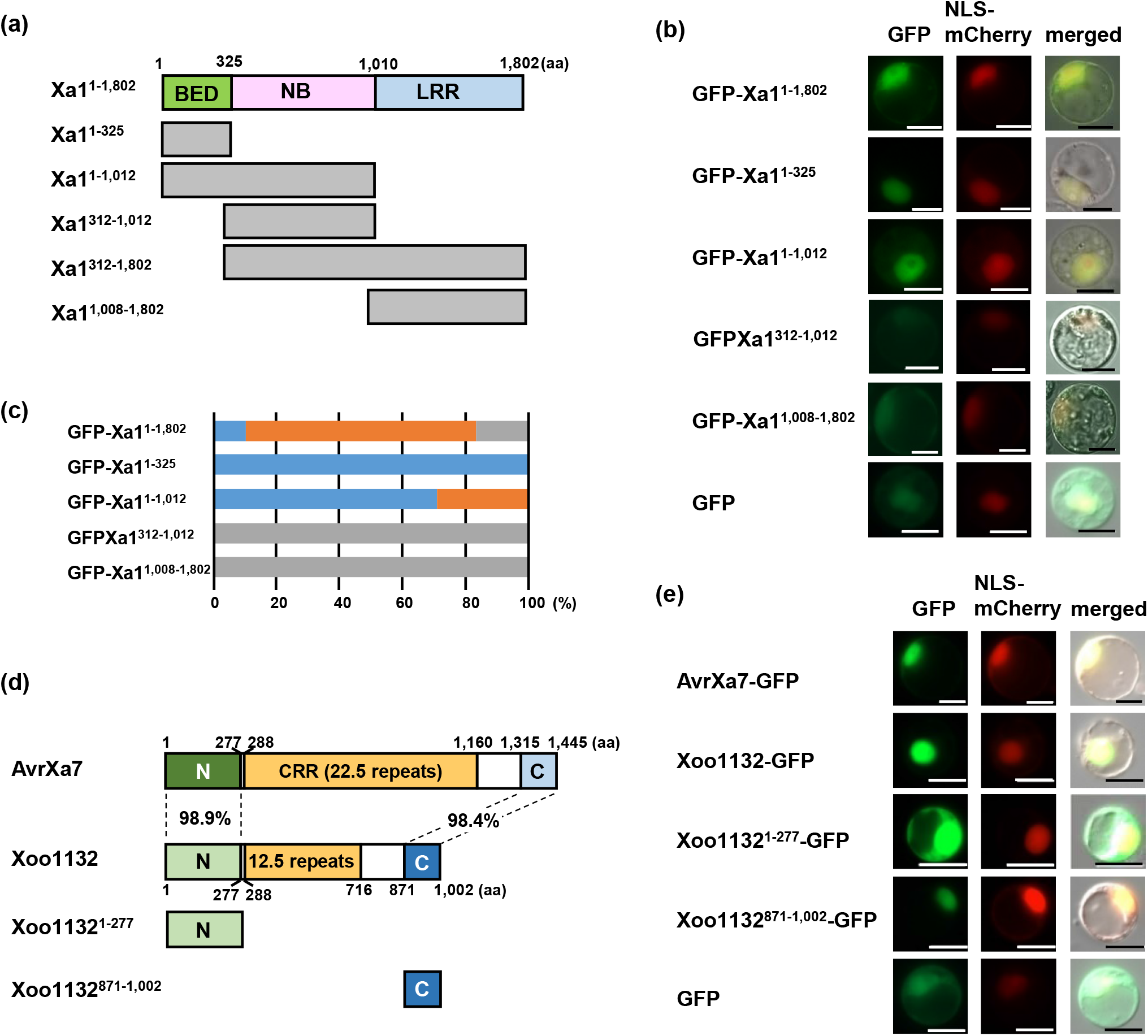
Subcellular localization of Xa1, AvrXa7, and Xoo1132. (a) Schematic structures of full-length Xa1^1-1,802^ and four truncated Xa1 fragments: Xa1^1-325^ (1–325 aa), Xa1^1-1,012^ (1–1,012 aa), Xa1^312-1,012^ (312–1,012 aa), and Xa1^1,008-1,802^ (1,008–1,802 aa). Numbers above the boxes indicate amino-acid residues. (b) Subcellular localization of the GFP-fused full length Xa1 and truncated Xa1 fragments in rice protoplasts. Fluorescence was observed 18 h after transformation. Scale bar, 10 μm. NLS-mCherry was used as a nuclear localization marker. (c) Proportions of cells showing nuclear and/or cytoplasmic localization of the Xa1 fragments. Blue and grey boxes indicate nuclear and cytoplasmic localization, respectively. Orange boxes indicate the cells in which fluorescence was detected in both the nuclei and cytoplasm. (d) Schematic structures of AvrXa7, Xoo1132, and two truncated Xoo1132 fragments. N, N-terminal region; CRR, central repeat region; C, C-terminal region. (e) Subcellular localization of the GFP-fused AvrXa7 and Xoo1132 and truncated Xoo1132 fragments in rice protoplasts. Fluorescence was observed 18 h after transformation. Scale bar, 10 μm. NLS-mCherry was used as a nuclear localization marker.

The *Xoo* strain T7174 (also called MAFF311018)(Ochiai *et al.*, 2005) is incompatible with the rice cultivar Kogyoku, which carries Xa1. *Xoo* T7174 contains 16 TAL effectors including AvrXa7 and Xoo1132 (Ochiai *et al.*, 2005). AvrXa7 and Xoo1132 have 98.9% sequence identity between their N-terminal regions and 98.4% identity between their C-terminal regions, and they have 22.5 and 12.5 repeats in their central repeat regions, respectively (Fig. 1d and Fig. S1). AvrXa7 and Xoo1132 were each fused to GFP and transiently expressed in rice protoplasts. Fluorescence from both constructs was detected in the nuclei (Fig. 1e). The N-terminal region (1–277 aa) and C-terminal region (871–1,002 aa) of Xoo1132 were also each fused to GFP. Xoo1132^871-1,002^-GFP was localized in the nucleus whereas Xoo1132^1-277^-GFP was detected in both the nucleus and the cytoplasm, as was also observed for the GFP control (Fig. 1e). This result was consistent with the fact that three nuclear localization signals (NLSs) exist at the C-terminal region of Xoo1132.

We next performed bi-molecular fluorescence complementation (BiFC) experiments to examine the interactions of Xa1 with AvrXa7 and Xoo1132. Full length Xa1^1-1,802^ protein was tagged with the C-terminal domain of Venus (Xa1^1-1,802^-Vc). Xoo1132 and AvrXa7 were each tagged with the N-terminal domain of Venus (Xoo1132-Vn and AvrXa7-Vn). However, when Xa1^1-1,802^-Vc was co-expressed with Xoo1132-Vn or AvrXa7-Vn in rice protoplasts, we failed to detect any fluorescence (Fig. 2a). Since it is possible that the tertiary structure of Xa1 might weaken the interaction with these TAL effectors, we tagged each of the Xa1 domains Xa1^1-325^, Xa1^312-1,012^, and Xa1^1,008-1,802^ with the N-terminal domain of Venus and used them in the BiFC experiments. Florescence was detected in the nuclei when Xa1^1-325^-Vc was co-expressed with Xoo1132-Vn or AvrXa7-Vn, but not when AvrXa7-Vn was co-expressed with Xa1^312-1,012^-Vc or Xa1^1,008-1,802^-Vc (Fig. 2a). These data suggest that the BED domain of Xa1 associates with these TAL effectors. We also co-expressed Xa1^1-325^-Vc with constructs containing Vn linked to either the N-terminal or C-terminal regions of Xoo1132 (Xoo11321-277 or Xoo1132^871-1,002^, respectively). The BED (Xa1^1-325^) domain interacted with Xoo1132^871-1,002^, but not with Xoo1132^1-277^ in the nucleus (Fig. 2a), even though the GFP constructs indicated that Xoo1132^1-277^ and Xoo1132^871-1,002^ are localized the nucleus (Fig. 1e).

**Fig. 2.**
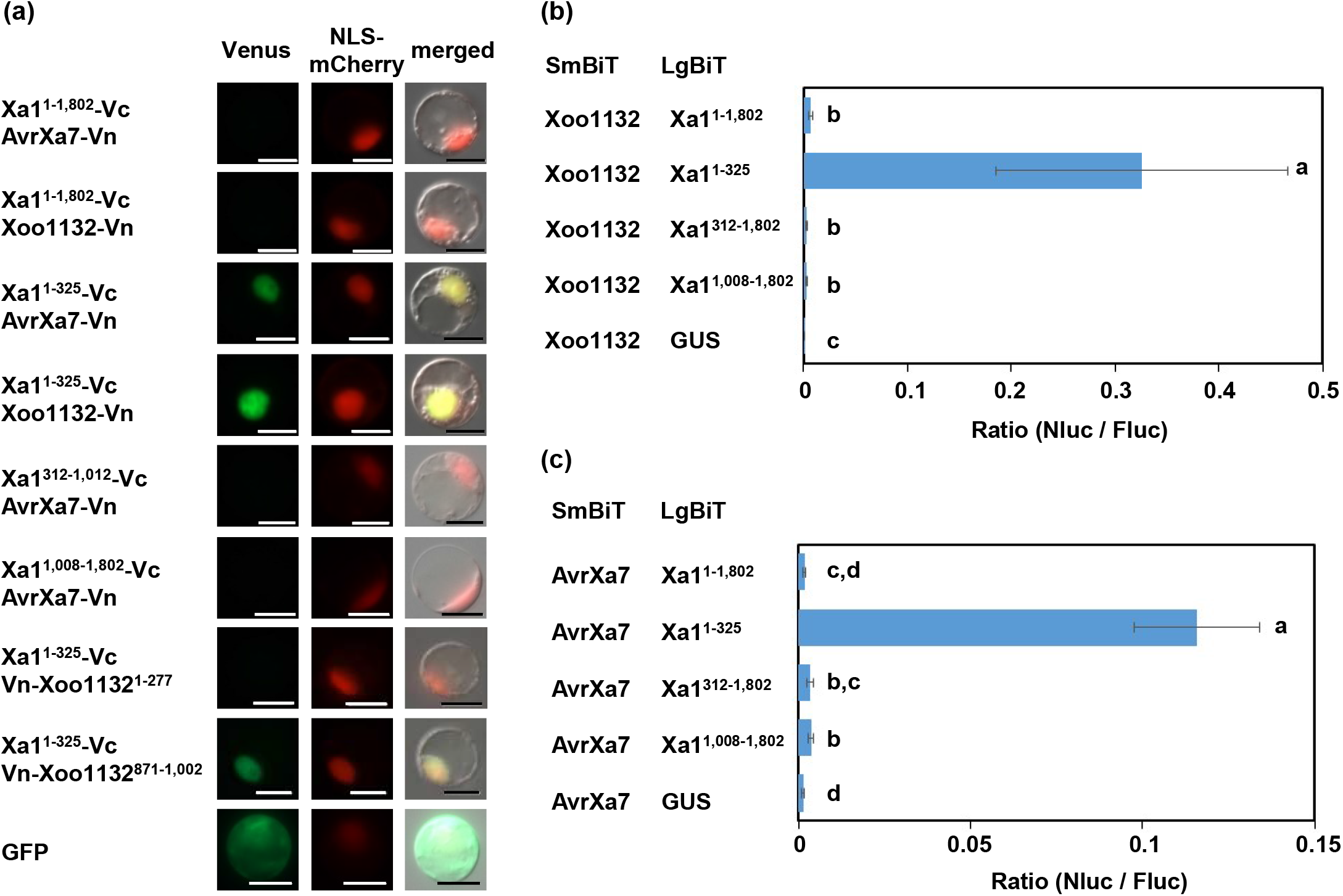
Xa1 interacts with AvrXa7 and Xoo1132. (a) Bimolecular fluorescence complementation (BiFC) analysis was used to visualize the interaction between the Xa1 fragments and the TAL effectors in rice protoplasts. NLS-mCherry was used as a nuclear localization marker. (b) Quantification of the interaction between the Xa1 fragments and Xoo1132. Split NanoLuc luciferase complementation assays were carried out by transient expression of SmBiT-fused Xoo1132 and LgBiT-fused Xa1 fragments. Values are means ± S.E. Different letters on the right sides of the data points indicate significant differences (*p* < 0.01). (c) Quantification of the interaction between the Xa1 fragments and AvrXa7. Split NanoLuc luciferase complementation assays were carried out by transient expression of SmBiT-fused AvrXa7 and LgBiT-fused Xa1 fragments. Values are means ± S.E. Different letters on the right sides of the data points indicate significant differences (*p* < 0.01).

To analyze the strength of the interaction between Xa1 and the TAL effectors, we carried out a split NanoLuc luciferase complementation assay (Taoka *et al.*, 2021). Xa1 and its fragments were fused to the Large BiT of NanoLuc Luciferase (LgBiT, 159 aa), and AvrXa7 and Xoo1132 were each fused to the Small BiT (SmBiT, 12 aa). The constructs were transfected into rice protoplasts with the *35S-Firefly Luciferase* (*Fluc*) construct, and the activities of both luciferases were measured using a microplate luminometer. Consistent with the BiFC experiments, Xa1^1-325^ (the BED domain) strongly interacted with both TAL effectors (Fig. 2b). Xa1^1-1,802^ interacted weakly but significantly with Xoo1132 when compared with the negative control (Fig. 2b). In addition, Xa1^312-1,802^ and Xa1^1,008-1,802^ displayed weak interaction with Xoo1132. AvrXa7 also exhibited similar interaction pattern as Xoo1132 (Fig. 2c). These data suggest that the BED domain of Xa1 may be the main region interacting with the TAL effectors, although the LRR domain may also be involved in the interaction.

The BiFC and split NanoLuc luciferase complementation experiments suggested direct interaction between the BED domain of Xa1 and the TAL effectors. To further test this possibility, we carried out yeast two-hybrid assays. However, interactions were not detected between the BED domain and either of the TAL effectors (Fig. S2). This result suggested that some host factor(s) might be required for the interaction between the BED domain and the TAL effectors.

### OsERF101 interacts with Xa1

To identify host factors involved in the Xa1-mediated immune response, we screened for Xa1 interactors using a yeast two-hybrid assay with the BED domain (Xa1^1-325^) and a rice cDNA library. Initially, we identified twelve candidates (Table S2). Among them, we selected the AP2/ERF type transcription factor OsERF101 /OsRAP2.6 (LOC_Os04g32620). Our selection was based on the predicted subcellular localization of OsERF101 and the reproducibility of its interaction with the BED domain. Yeast two-hybrid experiments indicated that OsERF101 interacted with the BED domain but not with other Xa1 domains (Fig. 3a).

**Fig. 3.**
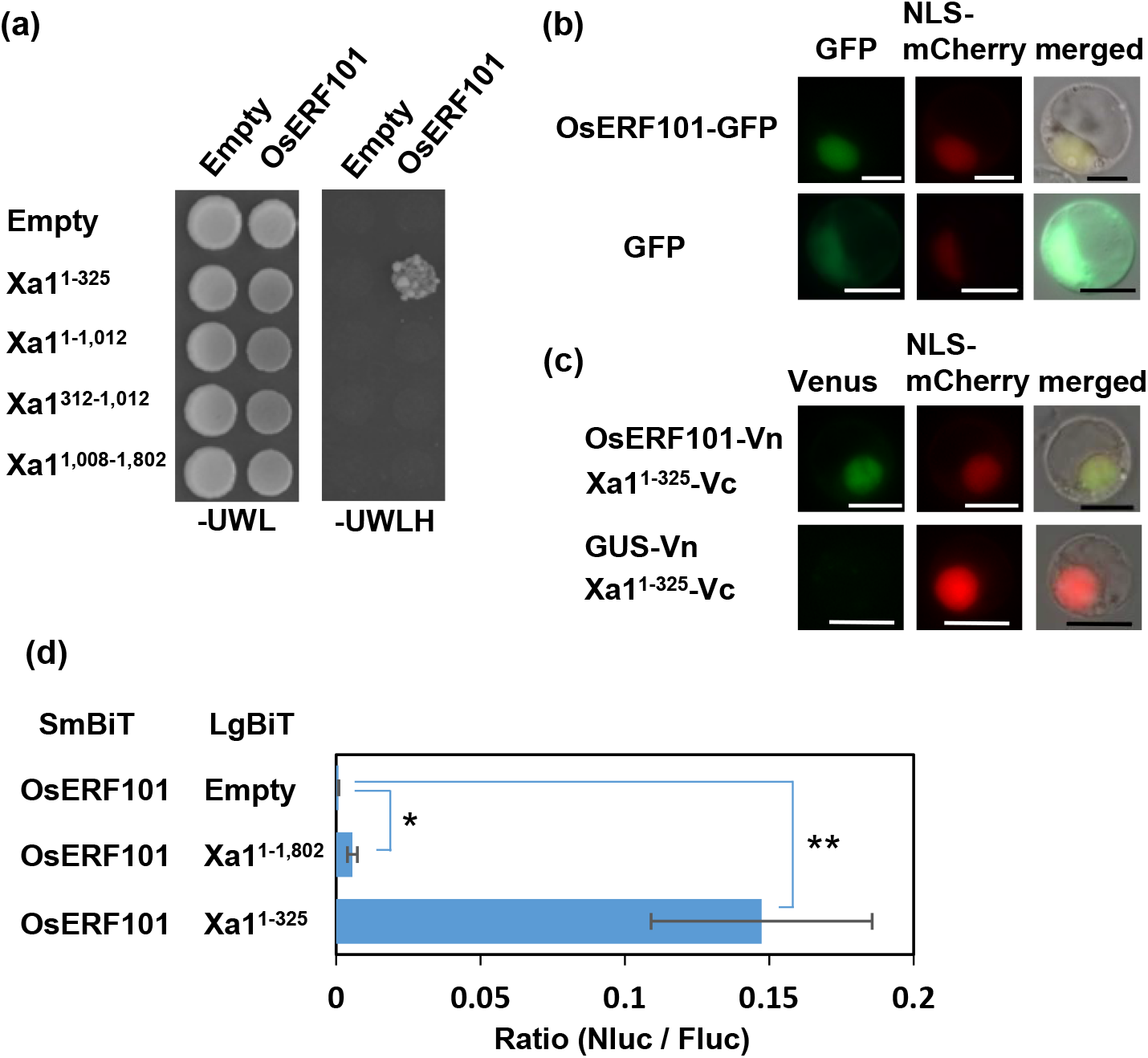
OsERF101 interacts with the BED domain of Xa1. Interaction between the Xa1 fragments and OsERF101 in yeast two-hybrid experiments. Growth of yeast colonies on –UWLH plates indicates a positive interaction. (b) Subcellular localization of GFP-fused OsERF101 in rice protoplasts. Fluorescence was observed 18 h after transformation. Scale bar, 10 μm. NLS-mCherry was used as a nuclear localization marker. (c) Bimolecular fluorescence complementation (BiFC) analysis was used to visualize the interaction between the Xa1 BED domain and OsERF101 in rice protoplasts. NLS-mCherry was used as a nuclear localization marker. GUS was used as a negative control. Scale bar, 10 μm. (d) Split NanoLuc luciferase complementation assays for quantification of the interaction between the Xa1 fragments and OsERF101. SmBiT-fused OsERF101 and LgBiT-fused Xa1 fragments were transiently expressed in rice protoplasts, and the luciferase activities were measured. Values are means ± S.E. Asterisks on the right sides of the data points indicate significant differences (*; *p* < 0.05; **; *p* < 0.01).

When the OsERF101 protein was fused with GFP (OsERF101-GFP) and transiently expressed in rice protoplasts, the protein was localized to the nucleus (Fig. 3b). We used BiFC assays to demonstrate *in planta* interactions between the BED domain and OsERF101 in the nucleus (Fig. 3c). In addition, we tested the interactions between Xa1 and OsERF101 using split NanoLuc luciferase complementation assays. We found that OsERF101 interacted strongly with the BED domain and weakly with the full length Xa1 protein (Fig. 3d).

### OsERF101 positively regulates bacterial blight resistance

We generated an *Xa1* knockout mutant (*xa1*) in the Kogyoku background using the CRISPR/Cas9 system. The mutation was caused by the generation of a stop codon created by a two base deletion (Fig. S3a). We then used a needleless syringe-infiltration technique to introduce suspensions of *Xoo* T7174 and *Xoo* T7133 (which is compatible with Kogyoku) into the leaves of Kogyoku plants. Wild-type plants developed HR lesions with dark brown edges and weak water soaking when infiltrated with *Xoo* T7174, but showed only water soaking when infiltrated with *Xoo* T7133 (Fig. 4a). In contrast, the *xa1* mutant did not develop HR lesions when infiltrated with *Xoo* T7174. Consistent with those results, the bacterial population of *Xoo* T7174 was much greater in the *xa1* mutant than in the wild-type plants (Fig. 4b). As mentioned in the Introduction, AvrXa7 induces *SWEET14* expression by direct binding to its promoter (Antony *et al.*, 2010). We observed stronger induction of *SWEET14* expression in the *xa1* mutant than in the wild-type plants after infiltration with *Xoo* T7174 (Fig. 4c).

**Fig. 4.**
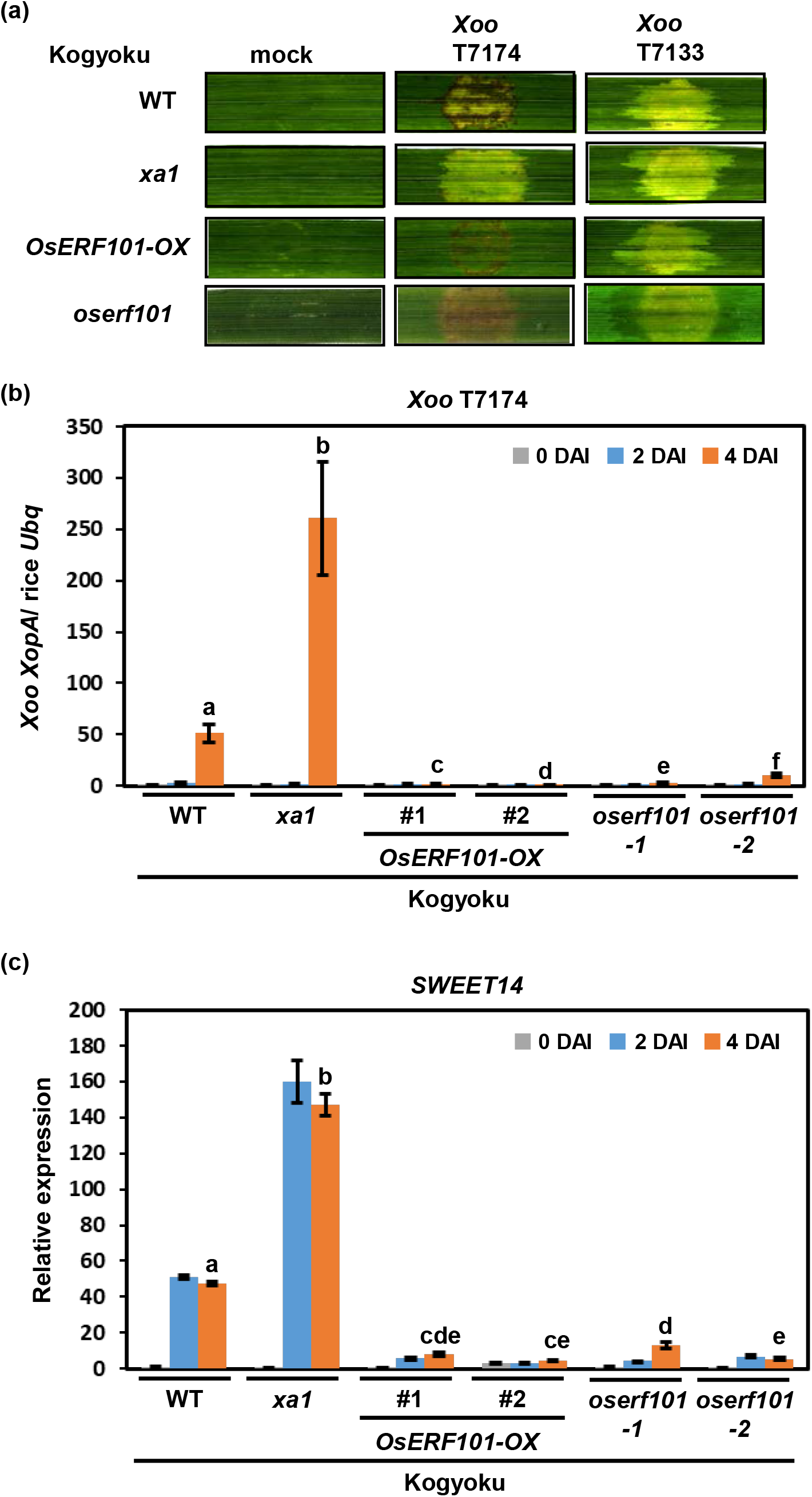
*OsERF101*-OX and *oserf101*enhance bacterial blight resistance in the Kogyoku background. (a) Hypersensitive response of the *OsERF101*-OX and *oserf101* plants in the Kogyoku background. The suspensions of *Xoo* T7174 or *Xoo* T7133 were injected into the leaves of three-week-old seedlings. Photos were taken at 4 days after inoculation (DAI). (b) The bacterial populations of *Xoo* T7174 in the *OsERF101*-OX and *oserf101* plants were analyzed by quantitative real-time PCR. The data indicate the DNA levels of the *X. oryzae XopA* gene relative to that of the rice *ubiquitin* gene. Values are means ± S.E. Different letters above the data points of 4 DAI indicate significant differences (*p* < 0.01). (c) Expression of *SWEET14* in the *OsERF101*-OX and *oserf101* plants after infection with *Xoo* T7174. The transcript levels were measured by quantitative real-time PCR. Values are means ± S.E. Different letters above the data points of 4 DAI indicate significant differences (*p* < 0.01).

To elucidate the function of OsERF101 in bacterial blight resistance, we generated transgenic Kogyoku plants overexpressing *OsERF101* using the CaMV *35S* promoter (Fig. S4). When these plants were infiltrated with the *Xoo* T7174 suspension, they exhibited a stronger HR than the wild-type plants, as indicated by the development of HR lesions without water soaking (Fig. 4a). Consistent with the stronger HR, both bacterial growth and *SWEET14* expression were reduced in the infiltrated *OsERF101*-OX plants when compared with the wild type (Fig. 4b,c). These results indicate that OsERF101 plays a positive role in resistance to bacterial blight.

### The knockout of *OsERF101* results in enhanced resistance

To analyze the involvement of OsERF101 in Xa1-mediated immunity, we used the CRISPR/Cas9 system to generate two knockout mutant lines of *OsERF101* in the Kogyoku background. The lines *oserf101-1* and *oserf101-2* both carried frame-shift mutations located approximately 220 bp from the start codon (Fig. S3b). When either of these lines were infiltrated with *Xoo* T7174, they exhibited light brown lesions that had a different appearance from the typical Xa1-induced HR lesion (Fig. 4a). This result was unexpected because OsERF101 is predicted to function as a positive regulator of rice immunity. However, the bacterial growth and *SWEET14* expression were strongly suppressed in the *oserf101* mutants after infiltration with T7174 (Fig. 4b,c), as we also observed in the *OsERF101*-OX plants. Thus, both the overexpression of *OsERF101* and the knockout of *OsERF101* induced strong resistance to rice bacterial blight.

### The enhanced resistance of the *OsERF101*-OX and *oserf101* plants depends upon *Xa1*

The *Xoo* strain T7133 is compatible with Kogyoku (Ogawa *et al.*, 1978) and produces disease lesions with water soaking when infiltrated into the leaves of Kogyoku (Fig. 4a). We determined the genome sequence of *Xoo* T7133 and found that it contains a type-A iTAL effector gene (Yoshihisa *et al.*, 2021). Thus far, all of the *Xoo* strains containing type-A iTAL effectors have been observed to inhibit Xa1-mediated immunity in rice (Ji *et al.*, 2020). Therefore, we expect that the iTAL effector of *Xoo* T7133 would also inhibit Xa1-mediated resistance.

We infiltrated a suspension of *Xoo* T7133 into the leaves of the *OsERF101*-OX plants. The *OsERF101*-OX leaves did not exhibit HR lesions as they did with *Xoo* T7174, but instead displayed water-soaked disease lesions (Fig. 4a). Consistent with this result, the bacterial populations of *Xoo* T7133 were increased in the *OsERF101*-OX plants, as they were in the wild-type plants and the *xa1* mutant (Fig. 5a). In addition, the expression levels of *SWEET14* in the *Xoo* T7133-infiltrated *OsERF101*-OX plants were the same as in the wild type and the *xa1* mutant (Fig. 5b). We also infiltrated a suspension of *Xoo* T7133 into the leaves of the *oserf101* knockout mutant lines. The atypical HR lesions induced by infection with *Xoo* T7174 were not observed after infection with *Xoo* T7133 (Fig. 4a). As observed in the *OsERF101*-OX plants, the infiltrated *oserf101* leaves developed water-soaked disease lesions (Fig. 4a), along with wild-type levels of bacterial growth and *SWEET14* expression (Fig. 5a,b). Because the iTAL effector of *Xoo* T7133 likely suppresses the immune responses induced via the recognition of TAL effectors by Xa1, these results suggest that the enhanced resistance of the *OsERF101*-OX and *oserf101* plants may be Xa1-dependent.

**Fig. 5.**
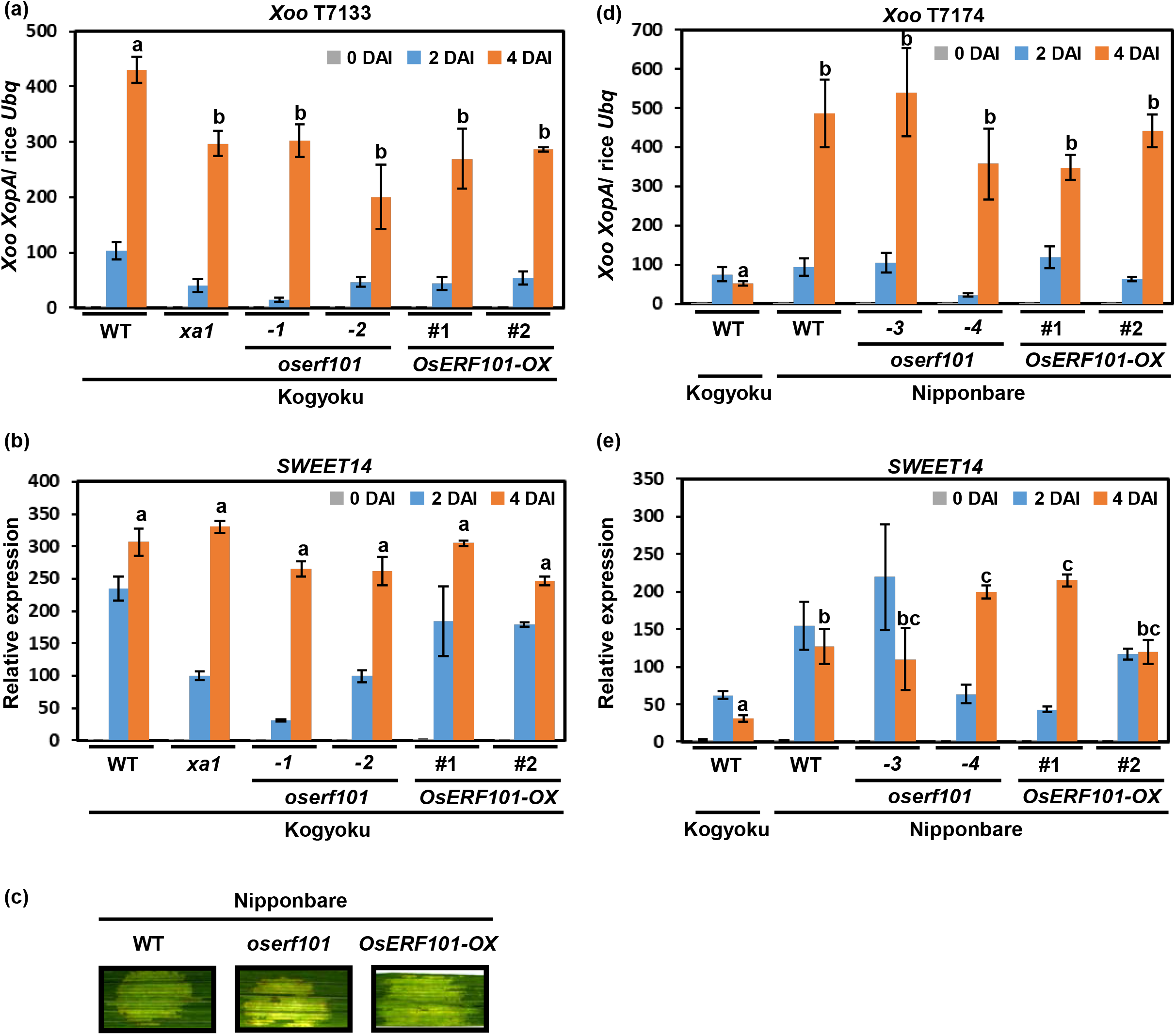
Enhanced resistance of the *OsERF101*-OX and *oserf101*plants depends on Xa1. (a) The bacterial populations of *Xoo* T7133 in the *OsERF101*-OX and *oserf101* plants in the Kogyoku background were analyzed by quantitative real-time PCR. The data indicate the DNA levels of the *X. oryzae XopA* gene relative to that of the rice *ubiquitin* gene. Values are means ± S.E. Different letters above the data points of 4 DAI indicate significant differences (*p* < 0.01). (b) Expression of *SWEET14* in the *OsERF101*-OX and *oserf101* plants after infection with *Xoo* T7133. The transcript levels were measured by quantitative real-time PCR. Values are means ± S.E. Different letters above the data points of 4 DAI indicate significant differences (*p* < 0.01). (c) Disease symptoms of the *OsERF101*-OX and *oserf101* plants in the Nipponbare background. The suspension of *Xoo* T7174 was injected into the leaves of three-week-old seedlings. Photos were taken at 4 DAI. (d) The bacterial populations of *Xoo* T7174 in the *OsERF101*-OX and *oserf101* plants in the Nipponbare background were analyzed by quantitative real-time PCR. The data indicate the DNA levels of the *X. oryzae XopA* gene relative to that of the rice *ubiquitin* gene. Values are means ± S.E. Different letters above the data points of 4 DAI indicate significant differences (*p* < 0.01). (e) Expression of *SWEET14* in the *OsERF101*-OX and *oserf101* plants in the Nipponbare background after infection with *Xoo* T7174. The transcript levels were measured by quantitative real-time PCR. Values are means ± S.E. Different letters above the data points of 4 DAI indicate significant differences (*p* < 0.01).

To further confirm that the enhanced resistance induced by overexpression and knockout mutations of *OsERF101* is dependent on Xa1, we generated transgenic plants overexpressing *OsERF101* and created knockout mutants of *OsERF101* in the rice cultivar Nipponbare background (Fig. S3c). Nipponbare does not possess the *Xa1* gene. The knockout mutant alleles of *OsERF101* in the Nipponbare background were named *oserf101-3* and *oserf101-4*. The *OsERF101-OX* and *oserf101* Nipponbare leaves developed water-soaked disease lesions when infiltrated with *Xoo* T7174 (Fig. 5c). In addition, we analyzed the bacterial growth and expression of *SWEET14* in the Nipponbare lines after infiltration with T7174 (Fig. 5d,e). The results indicated that neither overexpression of *OsERF101* nor the *oserf101* mutation caused enhanced resistance to *Xoo* T7174 in the Nipponbare background. Thus, it is likely that the enhanced resistance induced by both overexpression and knockout mutations of *OsERF101* in the Kogyoku background is dependent on Xa1.

### *OsERF101* overexpression and the *oserf101* knockout mutation induce different types of immune responses

We observed that the *OsERF101*-OX lines and the and *oserf101* knockout lines in the Kogyoku background displayed different types of HR lesions after infiltration with *Xoo* T7174 (Fig. 4a). This suggested that *OsERF101*-OX and *oserf101* may have different effects on the expression of downstream genes. Therefore, we carried out RNA-seq analyses using mRNAs purified from wild-type, *OsERF101*-OX, and *oserf101* leaves at 2 days after infiltration with *Xoo* T7174. We identified a total of 948 differentially regulated genes (false discovery rate < 0.1) whose expression levels in *OsERF101*-OX and/or *oserf101* were different from those in wild-type plants (Fig. 6a). Among the 381 genes that were upregulated in *OsERF101*-OX and/or *oserf101* when compared with the wild type, 327 were upregulated in either *OsERF101*-OX or *oserf101*, and the remaining 54 genes were upregulated in both plants. Similarly, the number of the genes down-regulated in both *OsERF101*-OX and *oserf101* was smaller than the number that were down-regulated in either *OsERF101*-OX or *oserf101* (Fig. 6a). Therefore, the overexpression of *OsERF101* affects the expression of a largely different set of downstream genes than those affected by the knockout mutation of *OsERF101*. Thus, it is likely that the mechanism by which the knockout of *OsERF101* enhances Xa1-mediated immunity is different from the mechanism by which *OsERF101* overexpression enhances immunity. These results suggest two regulatory pathways, both mediated by Xa1. In one pathway, OsEFR101 functions as a positive immune regulator, whereas the other pathway is negatively regulated by OsERF101.

**Fig. 6.**
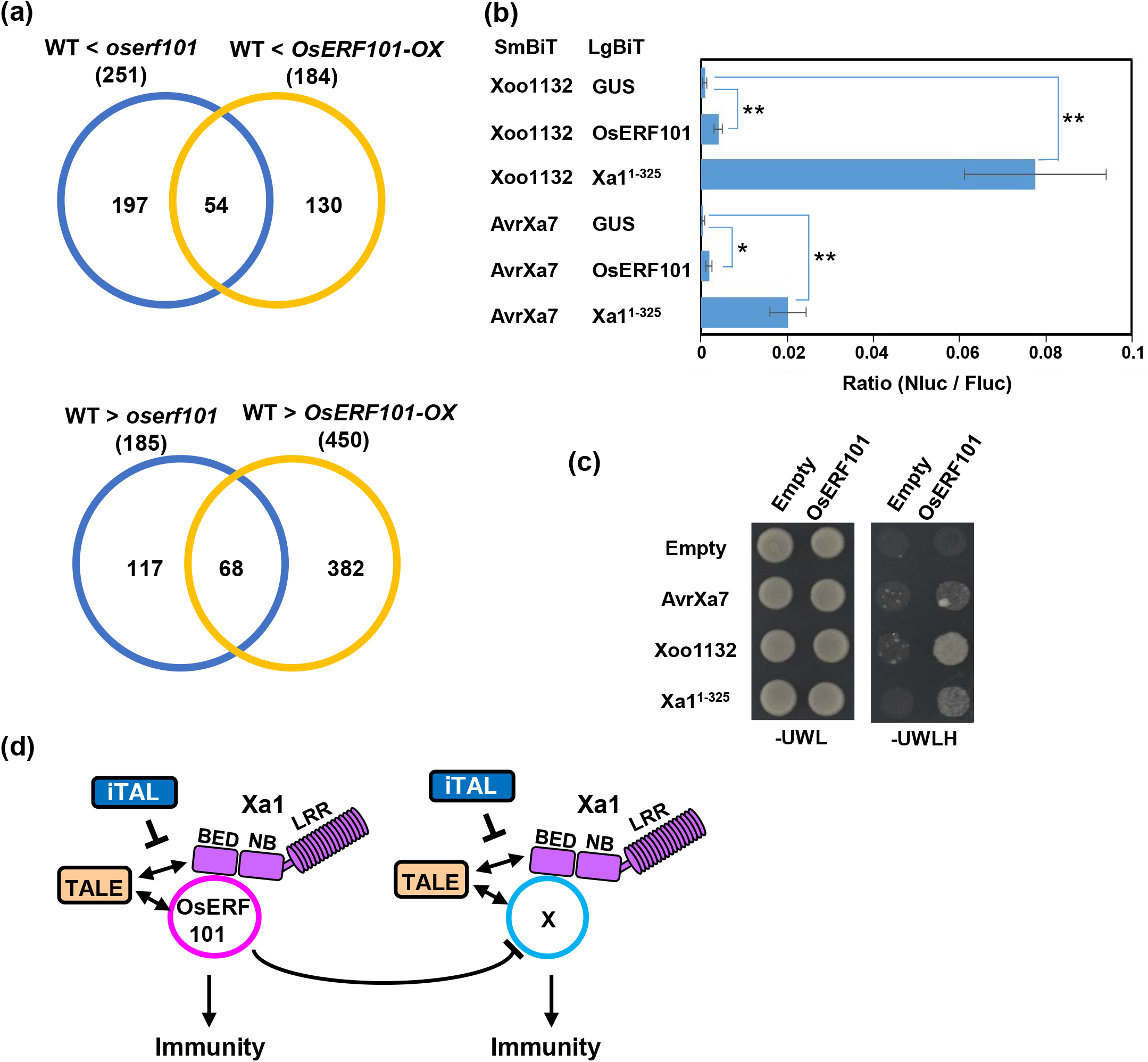
*OsERF101*-OX and *oserf101* plants activate different Xa1-mediated immunity pathways. (a) Comparative analysis of transcriptomes in wild-type, *OsERF101*-OX, and *oserf101* plants. RNA-seq analysis was carried out using leaves at 2 DAI with *Xoo* T7174. (b) Split NanoLuc luciferase complementation assays for quantification of the interaction between OsERF101 and the TAL effectors. SmBiT-fused AvrXa7 or Xoo1132 and LgBiT-fused ERF101 were transiently expressed in rice protoplasts, and the luciferase activities were measured. Values are means ± S.E. Asterisks on the right sides of the data points indicate significant differences (*, *p* < 0.05; **, *p* < 0.01). (c) Interaction between OsERF101 and TAL effectors in yeast two-hybrid experiments. Growth of yeast colonies on –UWLH plates indicates a positive interaction. (d) Proposed model for OsERF101-mediated immunity. OsERF101 interacts with Xa1, and positively regulates Xa1-mediated immunity. OsERF101 may negatively regulate a putative Factor X, which functions as a positive regulator of Xa1-mediated immunity. Depletion of OsERF101 may lead to enhance Factor X-mediated immunity.

### OsERF101 interacts with the TAL effectors

As described above, we did not detect direct interactions between the BED domain and the TAL effectors in yeast two-hybrid assays. This raised the possibility that OsERF101 may function as a link between Xa1 and the TAL effectors. Therefore, we looked for interactions between OsERF101 and the TAL effectors. The split NanoLuc luciferase complementation assays indicated that OsERF101 significantly interacted with the TAL effectors (Fig. 6b), although the interactions were much weaker than those between the BED domain and the TAL effectors. In addition, the yeast two-hybrid experiments demonstrated direct interactions between OsERF101 and the TAL effectors (Fig. 6c). Thus, it is possible that the recognition of the TAL effectors by Xa1 is dependent upon interactions with OsERF101.

## Discussion

Plants contain large numbers of NLR proteins. Some NLRs possess integrated decoy domains that are targeted by pathogen effectors. These integrated decoy domains mimic other effector targets whose binding to the effectors leads to disease development. Although BED was predicted to be an integrated decoy domain (Zuluaga *et al.*, 2017), direct evidence for this has not yet been obtained. Here, we showed that Xa1 forms complexes with the *Xoo* TAL effectors AvrXa7 and Xoo1132 via the BED domain, although direct interaction between the BED domain and the TAL effectors was not detected. We also found that the BED domain and the TAL effectors interact with the transcription factor OsERF101. Overexpression of *OsERF101* enhanced Xa1-mediated disease resistance, suggesting that OsERF101 plays a pivotal role in Xa1-mediated recognition of the TAL effectors and immune activation. Taken together, our results suggested that OsERF101 is a positive regulator in Xa1-mediated immunity.

On the other hand, we found that the *oserf101* knockout mutants also showed enhanced resistance to *Xoo* T7174. Interestingly, the HR lesions of the Kogyoku *oserf101* knockout mutants exhibited completely different characteristics from those of wild type and *OsERF101*-OX plants. Furthermore, the *OsERF101*-OX plants and the *oserf101* mutants influenced the transcription of largely different sets of downstream genes. This phenomenon suggests an additional Xa1-mediated immune pathway that is negatively regulated by OsERF101. Based upon these data, we propose a model for Xa1-OsERF101-mediated immune signaling by hypothesizing the existence of an “X factor” involved in Xa1-mediated immunity (Fig. 6d). In this model, both OsERF101 and the X factor positively regulate Xa1-mediated immunity, but OsERF101 has the ability to inhibit the activity of the X factor. Over-expression of OsERF101 enhances the Xa1-OsERF101-mediated immunity, but may suppress the X factor. The *oserf101* mutation results in enhancement of the X factor-mediated immune response. This model is consistent with two observations: 1) the enhanced resistance to *Xoo* T7174, induced by either the over-expression of *OsERF101* or the knockout of *OsERF101* in the Kogyoku background, is not observed with *Xoo* T7133, which carries an iTAL effector; 2) neither *OsERF101*-OX nor the *oserf101* mutation enhance *Xoo* T7174 resistance in the Nipponbare background, which does not carry Xa1. It should be noted here that *Xoo* T7174 also contains an iTAL effector named Tal3b, however, Tal3b is not functional in *Xoo* T7174 for unknown reasons (Ji *et al.*, 2016). We expect that the identification of the X factor will facilitate our understanding of Xa1-mediated immune signaling. Although we looked for rice genes up-regulated in the *oserf101* plants, we have not yet identified potential candidates for the X factor.

The *Xa1* alleles *Xa2*, *Xa14*, *Xa45*, and *Xo1* also encode BED-NLR proteins. The BED and NB domains are highly homologous among these Xa1 allelic members. However, they are differentiated by the numbers of repeats in their C-terminal LRR regions (Ji *et al.*, 2020; Read *et al.*, 2020; Zhang *et al.*, 2020), and they confer different resistance spectra to races of *Xoo*. Since the immune responses mediated by these allelic members are suppressed by iTAL effectors (Ji *et al.*, 2020), it is likely that the they all recognize the TAL effectors. The differences in the LRR regions suggest that the LRR domains may be the determinants of race specificity to *Xoo*. However, we observed only weak interactions between the LRR domain of Xa1 and the two TAL effectors in the split NanoLuc luciferase complementation assays. Instead, our BiFC and split NanoLuc luciferase complementation assays suggested the formation of a complex involving a TAL effector, OsERF101, and the BED domain of Xa1. Therefore, it is possible that the differences among the LRR regions of the Xa1 allelic members may affect either the formation of the tertiary complex or the LRR domain-mediated recognition of the tertiary complex.

The yeast two-hybrid experiments indicated that OsERF101 directly interacts with Xa1 and the TAL effectors. In addition, OsERF101 regulates Xa1-dependent immunity, and we speculate that the putative X factor also regulates Xa1-dependent immunity. Host factors that are targeted by pathogen effectors and act as co-receptors with NLRs are referred to as sensing decoys (Paulus & van der Hoorn, 2018). Therefore, it is possible that OsERF101 and the X factor may be sensing decoys targeted by the TAL effectors, and that they function as co-receptors with Xa1. Since sensing decoys generally promote effector recognition in the presence of their cognate NLR proteins (Paulus & van der Hoorn, 2018), this scenario is consistent with the fact that the over-expression and knockout mutation of *OsERF101* enhanced Xa1-dependent immunity in the Kogyoku background, but not in the Nipponbare background. However, if OsERF101 and the putative X factor function only as sensing decoys, the immune responses activated in the *OsERF101*-OX and *oserf101* plants should be same. The difference in the immune responses between the *OsERF101*-OX and *oserf101* plants imply that OsERF101 also contributes to the activation of Xa1-mediated immune signaling.

Consistent with recent reports (Read *et al.*, 2020; Xu *et al.*, 2021), we found that Xa1 is localized in the nucleus. Our data indicated that the BED domain can confer nuclear localization. Thus, it is likely that Xa1 recognizes the TAL effectors in the nucleus. In fact, the inhibition of Xa1-mediated immunity by the iTAL effectors requires the nuclear localization of the iTAL effectors (Ji *et al.*, 2016). These data strongly suggest that Xa1 activates immune responses within the nucleus. Recent investigations using co-immunoprecipitation indicated that the Xa1 allelic member Xo1 interacts with the iTAL effector Tal2h (Read *et al.*, 2020), although it is not yet known whether the interaction is direct or indirect. Although the molecular mechanisms of how the iTAL effectors inhibit Xa1-mediated immunity remain to be identified, it is possible that the iTAL effectors may suppress Xa1 through OsERF101 or the putative X factor.

It has been reported that the BED domains form a dimer (Zhang *et al.*, 2020). This suggests that oligomerization to form a resistosome may occur in BED NLRs, as occurs with other NLR proteins (Wang *et al.*, 2019; Ma *et al.*, 2020; Martin *et al.*, 2020). If interactions among the BED domain, OsERF101, and the TAL effectors alter the tertiary structure of the Xa1 protein, it is possible that this structural change may facilitate oligomerization through the BED domains. Recent structural studies of resistosomes using cryo-electron microscopy are beginning to reveal how NLR activates immunity (Wang *et al.*, 2019; Ma *et al.*, 2020; Martin *et al.*, 2020). Some NLRs with the N-terminal Toll-interleukin-1 receptor (TIR) domain have been reported to induce cell death through their nicotinamide adenine dinucleotide hydrolase activity (Horsefield *et al.*, 2019; Wan *et al.*, 2019). More recently, it was discovered that several NLRs with the N-terminal coiled coil domain function as plasma membrane-localized calcium-permeable channels (Bi *et al.*, 2021; Jacob *et al.*, 2021). However, the molecular mechanisms by which nuclear-localized NLRs activate immunity in plants are still unknown. This report on the interaction between Xa1 and OsERF101 in the initiation of the immune response provides new insight into NLR-mediated immunity.

## Acknowledgements

We thank Drs. Seiji Tsuge and Bing Yang for valuable suggestions concerning TAL effectors, Masaki Endo for the CRISPR/Cas9 expression constructs, Ken-ichiro Taoka for the split NanoLuc Luciferase assay constructs, Tetsuya Nakazaki and Kazusa Nishimura for use of their greenhouse, and Nao Hayata, Maho Izumitani, Shunsuke Ando and Toshikazu Ohuchi for technical assistance. This research was supported by Grants-in-Aid for Scientific Research (A)(19H00945), for Scientific Research on Innovative Areas (18H04789), for Exploratory Research (20K21320), Strategic International Collaborative Research project promoted by the Ministry of Agriculture, Forestry and Fisheries, Tokyo, Japan (JPJ0088379) and Basic Science Research Projects from the Mitsubishi Foundation to T.K.; by Grants-in-Aid for Scientific Research (JP15K18649) and Basic Science Research Projects from the Sumitomo Foundation to K. Y.

## Author contributions

SY and TK designed the research. AY, SY, SS, and KY performed the experiment. MS and AM analyzed RNAseq data. SY and TK wrote the manuscript. AY and SY contributed equally to this work.

## Figure legends

**Fig. S1.**
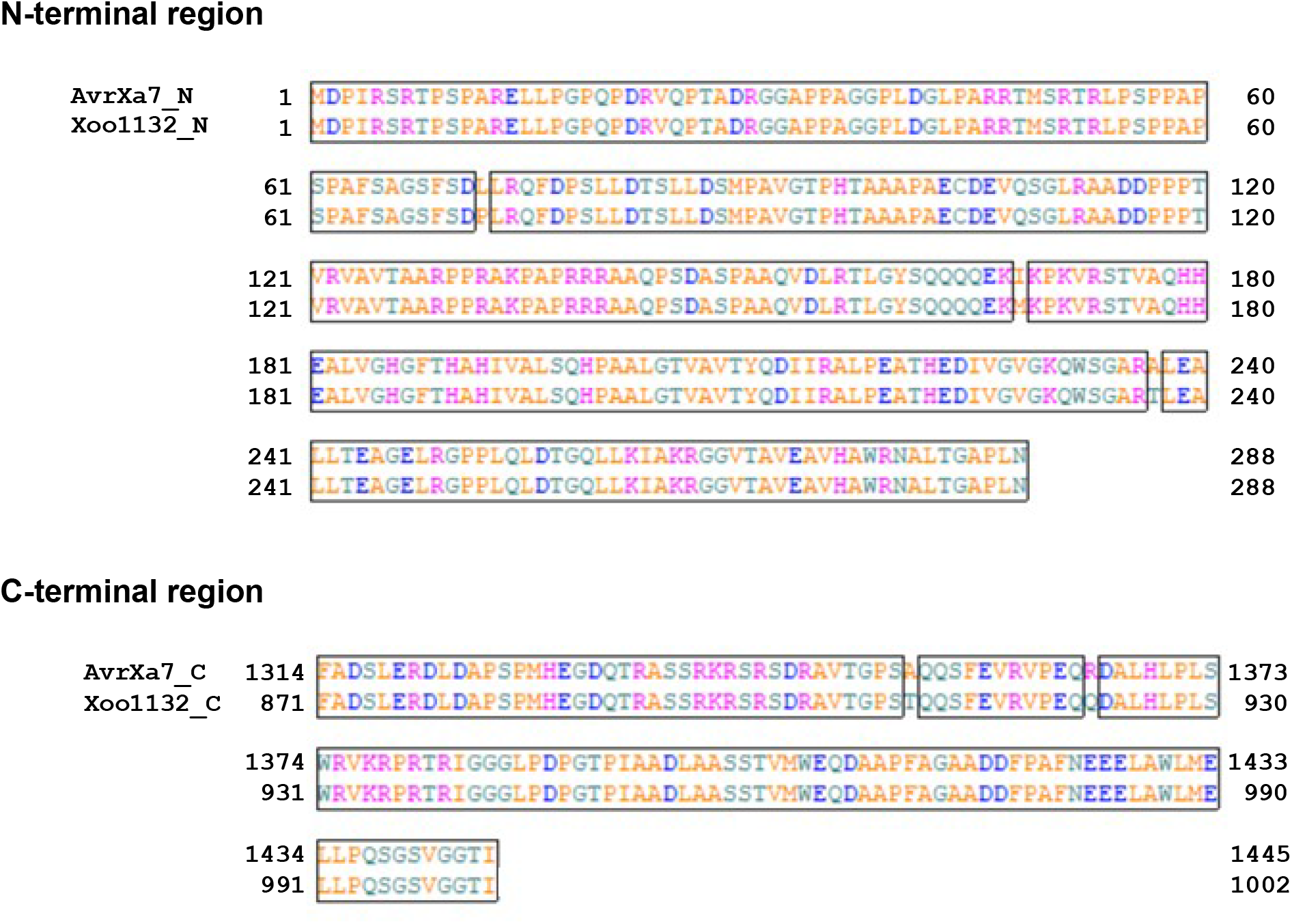
Alignment of the amino acid sequences of N- and C-terminal regions in Xoo1132 and AvrXa7.

**Fig. S2.**
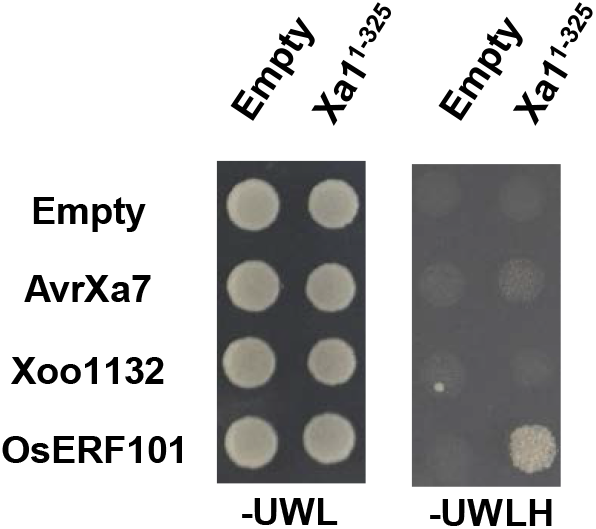
Interaction of Xa1^1-325^ with OsERF101, Xoo1132 or AvrXa7. Growth of yeast colonies on –UWLH plates indicates a positive interaction.

**Fig. S3.**
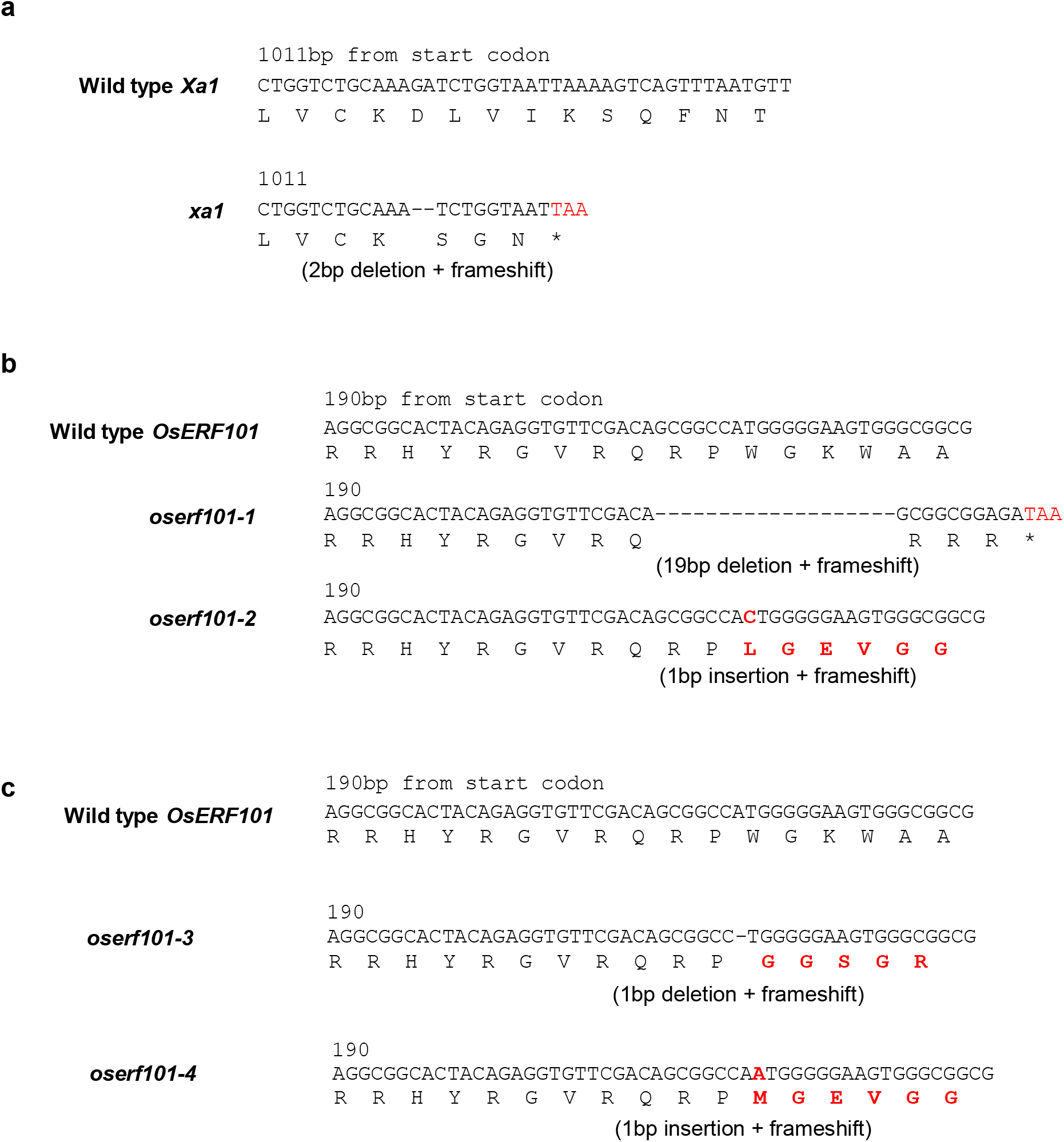
The mutants produced in this study. (A) The mutation site of *xa1*. (B) The mutation sites of *oserf101-1* and *oserf101-2* in the Kyogoku background. (C) The mutation sites of *oserf101-3* and *oeerf101-4* in the Nipponbare background.

**Fig. S4.**
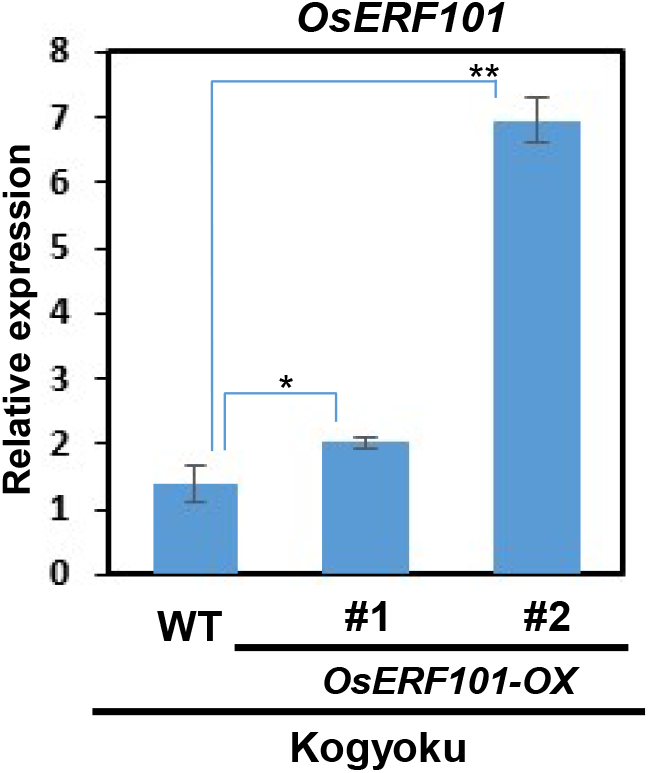
Transcript levels of *OsERF101* in the leaves of the transgenic plants overexpressing *OsERF101*. The transcript levels were measured by quantitative real-time PCR using specific primers of *OsERF101*. *Ubiquitin* was used as a control. Values are means ± S.E. Asterisks on the data indicate significant differences (*; p < 0.05; **; p < 0.01).

**Table S1.**
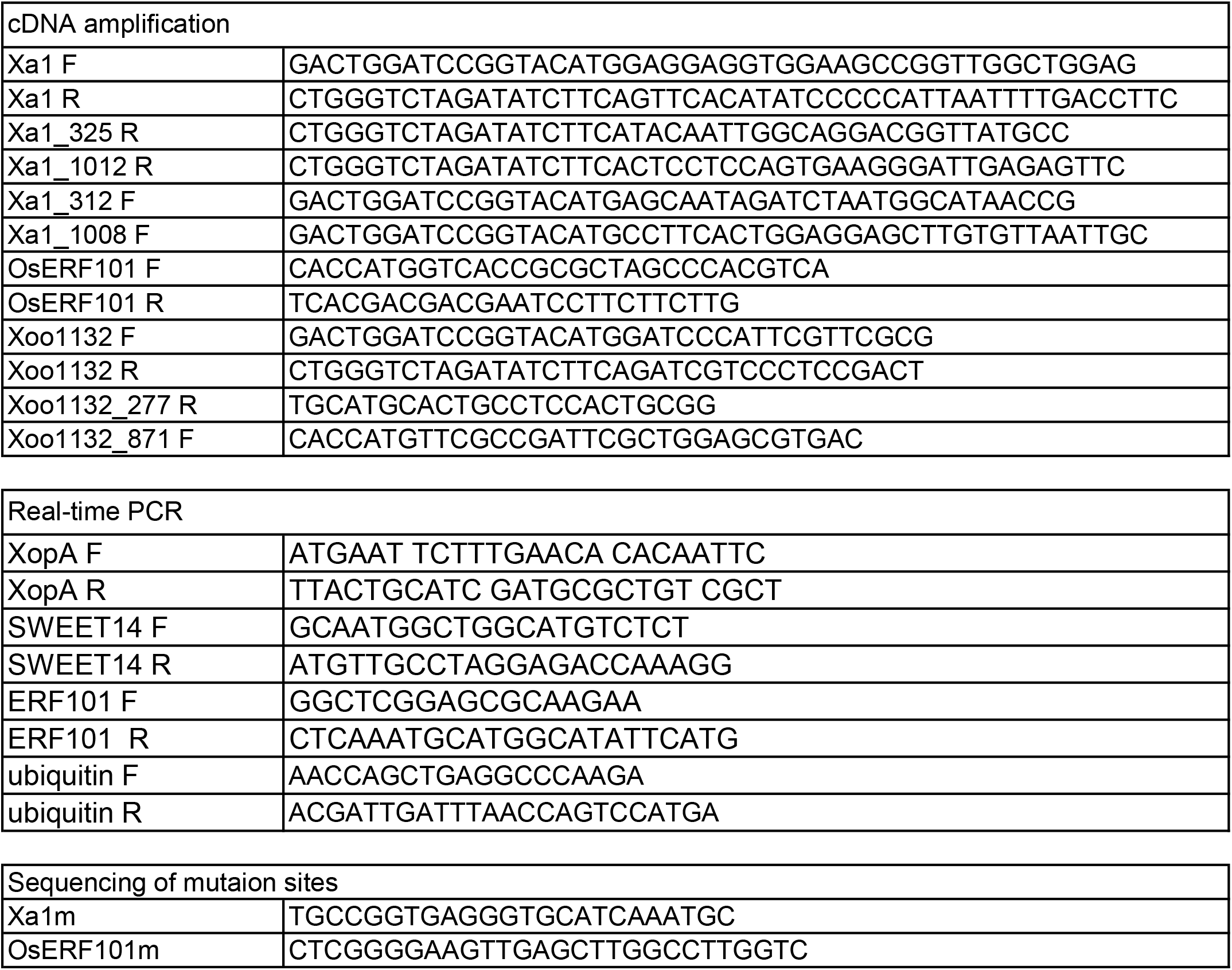
Primers used in this study

**Table S2.** The candidates of the Xa1 interactors.

| Gene ID | Description |
| --- | --- |
| Os02g0246300 | Prefoldin Domain |
| Os04g0398000 | Transcriptional factor(ERF) |
| Os01g0607200 | Amino acid permease protein |
| Os03g0215400 | Transcriptional factor(MADS box) |
| Os04g0610400 | Transcriptional factor(AP2) |
| Os01g0715900 | Pentatricopeptide repeat domain |
| Os03g0816700 | DUF567 family protein |
| Os05g0494500 | DUF250 domain |
| Os07g0438600 | Proteinase inhibitor I9 |
| Os08g0197400 | F-box domain |
| Os08g0477900 | Transcriptional factor (bHLH) |
| Os11g0106400 | Ubiquitin-activating enzyme E1 2 |

